# Protein-based cell population discovery and annotation for CITE-seq data identifies cellular phenotypes associated with critical COVID-19 severity

**DOI:** 10.1101/2024.03.14.584720

**Authors:** Denise Allen, Matthew Weaver, Sam Prokopchuk, Fritz Lekschas, Mike Jiang, Greg Finak, Evan Greene, Andrew McDavid

## Abstract

Technologies such as Cellular Indexing of Transcriptomes and Epitopes sequencing (CITE-seq) and RNA Expression and Protein sequencing (REAP-seq) augment unimodal single-cell RNA sequencing (scRNA-seq) by simultaneously measuring expression of cell-surface proteins using antibody derived oligonucleotide tags (ADT). These protocols have been increasingly used to resolve cellular populations that are difficult to infer from gene expression alone, and to interrogate the relationship between gene and protein expression at a single-cell level. However, the ADT-based protein expression component of these assays remains widely underutilized as a primary tool to discover and annotate cell populations, in contrast to flow cytometry which has used surface protein expression in this fashion for decades. Therefore, we hypothesized that computational tools used for flow cytometry data analysis could be harnessed and scaled to analyze ADT data. Here we apply Ozette Discovery™, a recently-developed method for flow cytometry analysis, to re-analyze a large (>400,000 cells) published COVID-19 CITE-seq dataset. Using the protein expression data alone, Ozette Discovery is able to identify granular, robust, and interpretable cellular phenotypes in a high-throughput manner. In particular, we identify a population of CLEC12A+CD11b+CD14- myeloid cells that are specifically expanded in patients with critical COVID-19, and can only be resolved by their protein expression profiles. Using the longitudinal gene expression data from this dataset, we find that early expression of interferon response genes precedes the expansion of this subset, and that early expression of PRF1 and GZMB within specific Ozette Discovery phenotypes provides a RNA biomarker of critical COVID-19. In summary, Ozette Discovery demonstrates that taking a protein-centric approach to cell phenotype annotation in CITE-seq data can achieve the potential that dual RNA/protein assays provide in mixed samples: instantaneous *in silico* flow sorting, and unbiased RNA-seq profiling.

**HIGHLIGHTS:**

- Ozette Discovery provides an alternative method for data-driven annotation of granular and homogeneous cell phenotypes in CITE-seq data using protein expression data alone.
- Our approach inherently accommodates for batch effects, and our novel background-normalization method improves the signal:noise ratio of these notoriously noisy protein measurements.
- While these subpopulations are not derived from RNA profiles, they have distinct and interpretable RNA signatures.
- We find a population of CLEC12A+CD11b+CD14- myeloid cells associated with critical COVID-19 severity that can only be identified by their protein profiles, and identify early expression of interferon response genes in a CD4 T cell subset as a predictor of CLEC12A+CD11b+CD14- cell expansion.
- Peforming differential expression analysis within our identified phenotypes reveals predictors of COVID-19 severity that are not found with coarser annotations.

## INTRODUCTION

Single-cell assays such as flow cytometry and single-cell RNA sequencing (scRNA-seq) quantify the abundance of protein and mRNA, respectively, and have each been critical to answering fundamental questions across biology. Multi-omic methods such as Cellular Indexing of Transcriptomes and Epitopes sequencing (CITE-seq)^1^ and RNA Expression and Protein sequencing (REAP-seq)^2^ seek to combine the advantages of flow cytometry and scRNA-seq by measuring mRNA levels in addition to expression of cell surface proteins using antibodies conjugated to oligonucleotide DNA barcodes. These techniques are now being used to disentangle the complex interplay between gene and protein expression in preclinical work, clinical trials, and large prospective cohorts^3,4^, including for the etiology of emerging diseases such as COVID-19^5^.

A critical step in the analysis of single-cell assays is identifying groups of cells that share a common cellular phenotype. In unimodal scRNA-seq, the most common methods take an unsupervised approach, based on reducing the dimension of the gene expression vector (e.g. via principal component analysis) and then detecting communities of cells in this reduced space^6^. These methods have the ability to discover cellular phenotypes *de novo*, but inherit the disadvantage that even well-known phenotypes require annotation in some fashion, by e.g. examining lists of genes that are differentially expressed between clusters. To address this issue, supervised methods to cluster the expression vectors and annotate cellular phenotypes have also been proposed^7,8^. These have the advantage of avoiding laborious manual annotation of clusters, but the phenotypes one can discover are limited by the coverage and quality of the reference dataset used to train the annotation model. Moreover, the process of recovering clusters not present in the reference, if possible at all, is not always reliable. Supervised and unsupervised methods both also often require extensive batch correction prior to cell phenotype annotation to ensure that the reduced features, and therefore cell clustering, are not unduly influenced by technical variation between batches.

Most clustering approaches for multi-omic data resemble the approaches used in unimodal scRNA-seq, attempting to blend protein and mRNA data in some beneficial way to yield a reduced dimensional vector that reflects expression in both modalities^9^. Although appealing from a statistical standpoint, blending methods often sacrifice interpretability, as the clusters produced reflect an amorphous combination of gene and protein expression. Furthermore, like unimodal scRNA-seq approaches, these methods often first require complex batch correction methods. Some methods have been developed that attempt to primarily or solely use the antibody-derived tag (ADT) protein data, but they are less common. scGate^10^ performs “hierarchical gating” on cells based on their ADT expression, sequentially examining one or two markers at a time to partition the protein expression space into discrete cell types. However, this method requires manual tuning per-marker to establish positive and negative thresholds, and is limited to closed-reference classification of predefined cell types. SECANT^11^ emphasizes supervised classification using previously-generated reference sets defined from ADT data, and similarly to scGate, requires heuristics to identify *de novo* cell types not present in the reference. CITE-sort^12^ is an unsupervised method that clusters cells by fitting parametric statistical models to a sequence of lower-dimensional projections of the ADT data. Although the authors illustrated the CITE-sort method on a dataset of 16,000 cells to discover 12 phenotypes, it does not appear to have been applied on additional larger-scale or higher-dimensional datasets since publication. In reviewing these currently-available methods, we therefore conclude that the multi-omic sequencing field is still lacking methods that can discover novel cell phenotypes in large datasets in a human-interpretable manner.

In flow cytometry, although unsupervised machine learning tools have been developed^13,14^, the gold-standard method of analysis remains manual hierarchical gating. Hierarchical gating maintains its appeal due to its simplicity and interpretability, but is limited by the throughput and lack of reproducibility between human operators. However, a recently-published method by Greene et al.^15^ proposes an approach that rapidly and exhaustively annotates cell phenotypes in single-cell protein data using the basic principles of hierarchical gating, while overcoming the current limitations in throughput and human intervention. This method has already been used to identify novel immune cell phenotypes in diverse disease contexts, including regulatory T cell (Treg) phenotypes in COVID-19^16^ and head and neck squamous cell carcinomas^17^. Ozette Technologies, a life science technology company specializing in the analysis of single-cell data, has developed the Ozette Discovery™ platform to build upon the methodology described in Greene et al. to provide a high-throughput and high-accuracy platform for the annotation of cytometry data. Since both flow cytometry and ADT data are antibody-based measures of protein expression, we hypothesized that Ozette Discovery could also be applied to ADT data to annotate cell phenotypes in single-cell sequencing data. If successful, this approach would enable exhaustive annotation of multi-omic data without use of predetermined reference datasets, and the protein-centric annotations would better harmonize with known cytometry phenotypes.

Here we demonstrate the application of Ozette Discovery to re-analyze a large (>400,000 cells) CITE-seq dataset of peripheral blood mononuclear cells (PBMCs) from COVID-19 patients and healthy controls. In the original publication of these data^5^, the authors, Liu et al., primarily identified cellular phenotypes using unsupervised Louvain clustering on the ADT data. These clusters, initially only identified by an integer, were annotated into biologically-meaningful phenotypes by examination of differentially expressed markers between clusters. In some cases this unsupervised approach was not sufficient, and the authors additionally had to perform manual gating in order to identify rare or complex cellular populations. This approach reflects the state-of-the-art, yet only enabled identification of 30 phenotypes across their ∼400,000 cells. In contrast, we find that Ozette Discovery, which also uses the protein expression data alone, identifies over 100 phenotypes that are more homogeneous and interpretable than those identified in the original publication. We also demonstrate how this homogeneity within each phenotype is important for accurately identifying cell populations whose abundance and gene expression profiles correlate with disease severity. Together, this work demonstrates that protein-centric cell annotation of CITE-seq data using Ozette Discovery enables significant progress towards achieving instantaneous flow sorting and RNA-seq profiling of mixed samples.

## RESULTS

### The Ozette Discovery™ method

To demonstrate the application of Ozette Discovery to CITE-seq ADT data, we re-analyzed the dataset presented in Liu et al., 2021^5^. This paper used CITE-seq to profile PBMCs from healthy donors and patients with moderate, severe, or critical COVID-19 at various points after symptom onset in order to identify cellular hallmarks of COVID-19 severity and how these hallmarks change over time (Fig 1A). This is one of the largest CITE-seq datasets publicly available in terms of the number of subjects included (47), proteins profiled (188), and number of cells sequenced (∼400,000). We started with the authors’ unfiltered feature by barcode matrices (GEO: GSE161918) and the results of their genotype-based donor demultiplexing scheme, which was combined with sample multiplexing via hashtag oligos. We identified cell-containing droplets^18,19^, demultiplexed samples using the genotype information and hashtag oligos^20^, and detected doublet barcodes^21,22^ using publicly-available methods, and then applied common quality control (QC) metrics (Fig 1B, methods).

**Figure 1:**
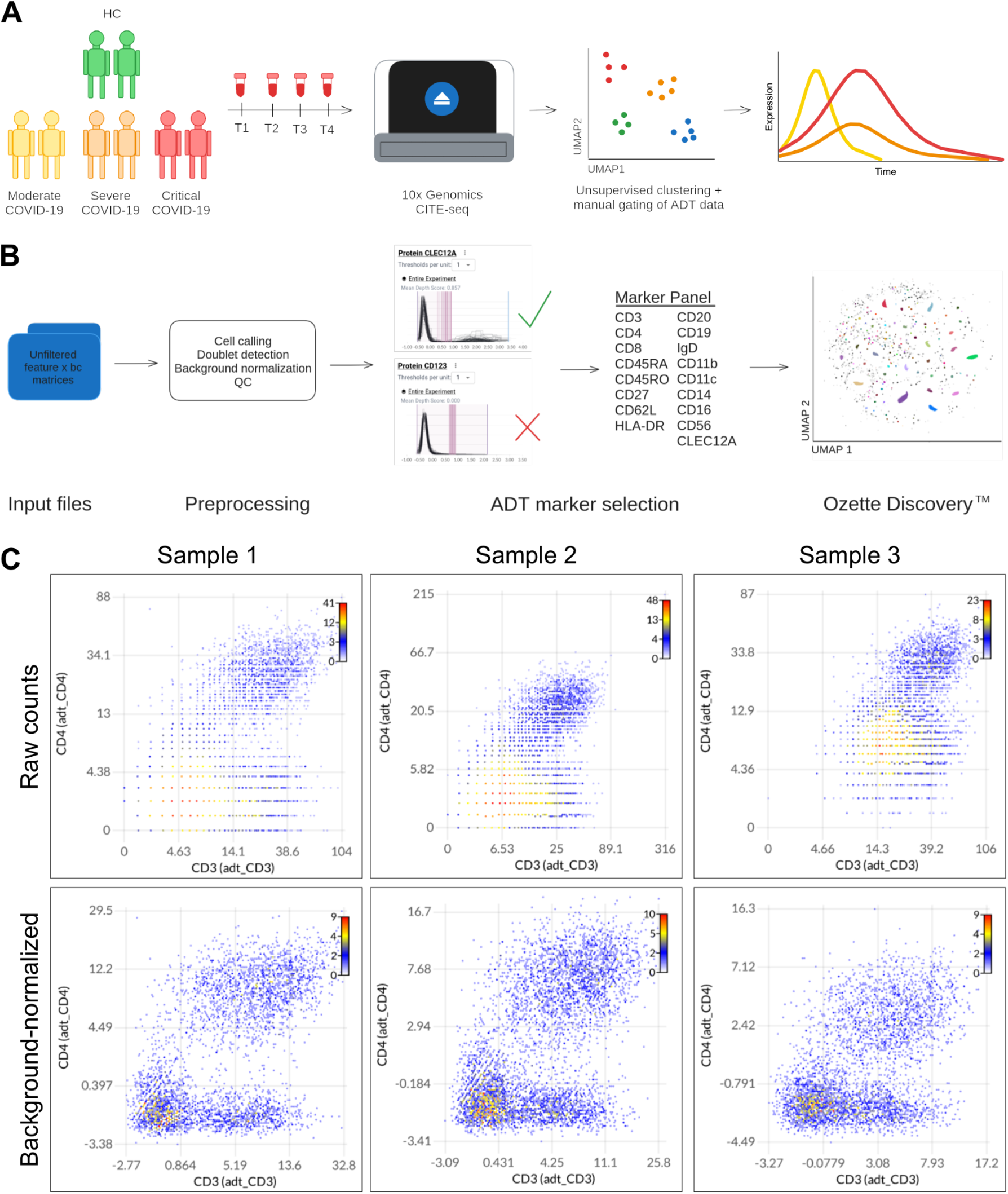
Applying Ozette Discovery to CITE-seq ADT data. **A)** Schematic of experimental design presented in Liu et al. 2021. PBMC samples were collected from healthy control (HC) subjects and patients with moderate, severe, or critical COVID-19 at various time points after symptom onset. These samples were profiled by CITE-seq using the 10x Genomics platform in order to identify expression patterns associated with COVID-19 severity over time. **B)** Schematic of Ozette Discovery analysis of Liu et al. 2021 CITE-seq ADT data. The inputs to the analysis were the authors’ unfiltered feature barcode matrices, on which cell calling, doublet detection, background normalization, and quality control were performed. Cell lineage or state markers and highly variable markers (e.g. CLEC12A) were prioritized whereas poorly staining or non-varying markers (e.g. CD123) were discarded, resulting in a panel of 17 markers considered. Finally, we used Ozette Discovery to annotate cell phenotypes in the dataset based on expression of the selected markers. Results were visualized in a UMAP where each point represents a cell and colors indicate distinct cell phenotypes. **C)** 2-D histograms showing expression of CD3 vs CD4 by CITE-seq ADT in three representative samples (columns). Top row shows distribution of cells based on raw ADT counts of UMIs. Bottom row shows the distribution of cells based on background-normalized ADT counts of UMIs. Axes are arcsinh transformed with cofactor = 7. Color scheme indicates the density of cells at each point (scale at top right of each plot).

CITE-seq data are notorious for having high levels of nonspecific background, which is thought to arise largely from unbound antibody tags that are not adequately washed away after staining. Since flow cytometry clustering methods–including Ozette Discovery–generally assume that a clear expression threshold exists between cells that do and do not express each protein measured, this ambient contribution makes it difficult to resolve true positive versus negative populations^23^. Ideally, there should be a large margin between positive and negative events, such that a threshold can be identified that is sensitive and specific. We found that existing normalization approaches for CITE-seq data^23^ did not sufficiently improve this margin between positive and negative populations, so we developed a generalized linear model (GLM)-based background normalization method (see Methods: Background Normalization) that materially improves the signal-to-noise ratio of CITE-seq ADT data (Fig 1C).

Even after this background normalization, not all markers had reliable thresholds between positive and negative events due to limited antibody avidity or small numbers of positive cells. Therefore, we prioritized markers that are well established PBMC lineage markers or highly variable across the dataset to input into the Ozette Discovery algorithm (Fig 1B). We first selected major cell lineage markers CD3, CD4, CD8, CD19, CD20, IgD, CD56, CD11b, CD11c, CD14, and CD16, as well as differentiation markers including CD45RA, CD45RO, CD27, HLA-DR, and CD62L. We also included the most variable marker across the dataset, CLEC12A (also known as CLL1). CLEC12A is a C-type lectin domain family receptor known to be a negative regulator of granulocyte and monocyte function, and has been reported as a biomarker of COVID-19 survival^24^. Ozette’s Discovery algorithm determines the threshold position between positive and negative expression for each of the selected markers, and then partitions the cells in the dataset into unique phenotypes based on where they fall relative to these thresholds and how frequently they are detected across samples^15^ (Fig 1B). Downstream analyses can then determine how the expansion, contraction, or gene expression patterns of these annotated cellular phenotypes are associated with biological variables of interest (Fig 5-Fig 7).

### Phenotypes identified by Ozette Discovery are granular, robust, and human-interpretable

As described above, Liu and colleagues partitioned their dataset into 30 phenotypes using a combination of unsupervised clustering using Seurat v3^25^ and manual gating in FlowJo. Executing this kind of manual gating followed by using lists of differentially expressed markers to map numbered clusters to meaningful cell types requires expert subject knowledge and a significant time investment. In contrast, application of Ozette Discovery to the CITE-seq ADT data presented in Liu et al. identified 170 phenotypes that are explicitly labeled by their ADT expression profile (Fig 2A, representative phenotypes labeled). These phenotypes can be visualized on the Ozette Discovery platform using an annotation-transformation UMAP^15^ (Fig 2A), which normalizes the marker expression and aligns cells from different samples so that each phenotype localizes together in the UMAP embedding. The Ozette Discovery platform also provides “backgating plot” visualizations to enable the direct interrogation of how each subpopulation was identified (Fig 2B). Backgating plots are one- or two-dimensional histogram plots showing the expression of one or two markers across all cells in a dataset, overlaid with the expression data for a particular subpopulation and the determined thresholds between positive and negative populations. In contrast to parsing through lists of overlapping differentially expressed markers between clusters, we can use these backgating plots to easily visualize and interrogate the basis of Ozette Discovery’s labels: the computationally-determined thresholds between positive and negative populations (Fig 2B, light blue lines) can be used to verify that the cells in each labeled population (Fig 2B, dark blue) fall confidently above or below these thresholds relative to all cells in the dataset (Fig 2B, gray).

**Figure 2:**
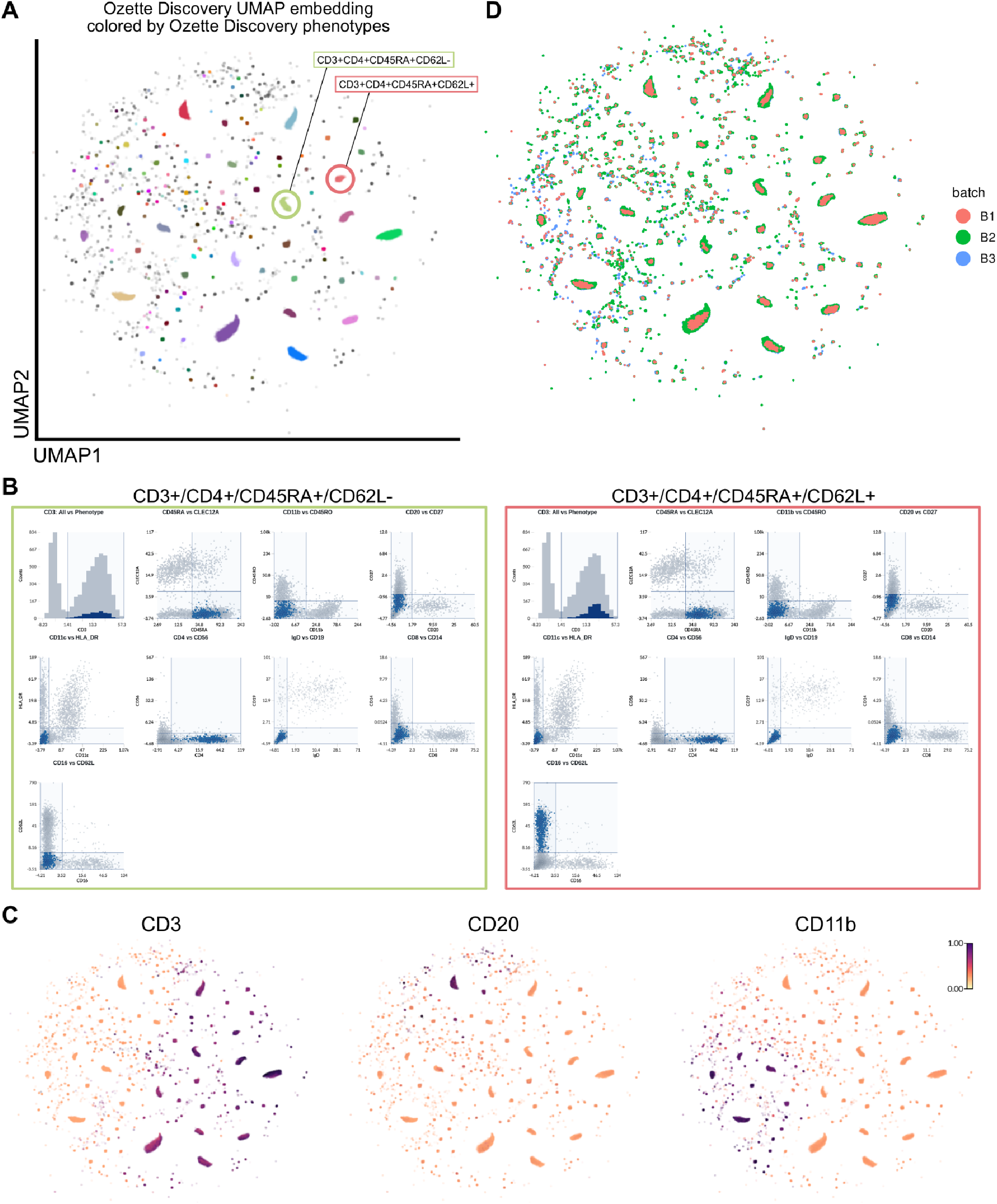
Ozette Discovery rapidly identifies granular, robust, and interpretable cell phenotypes from ADT data. **A)** Annotation transformation-based UMAP embedding showing Liu et al. data as annotated by Ozette Discovery. Clusters are colored by phenotype labels. Phenotypes expressing CD3+CD45RO+CD27+CD4+ or CD3+CD45RO+CD27+CD4+CD62L+ and negative for all other markers are circled and labeled. These phenotypes are also highlighted in (B/D). **B)** Series of backgating plots generated by Ozette’s Discovery platform demonstrating the distribution of all cells (gray) compared to the distribution of a selected phenotype (dark blue), relative to the computationally-determined thresholds between negative and positive populations for each marker (light blue lines). The color outlining each backgating plot series matches the color circling the corresponding phenotype in (A). Axes are arcsinh transformed with cofactor = 7. **C)** The same UMAP as in (A) colored by normalized ADT expression of CD3 (left), CD20 (middle), or CD11b (right). Color scale on the far right is the same for all plots and shows the relative expression from white (no expression) over red (medium expression) to dark purple (highest expression). **D)** The same UMAP as in (A) colored by the experimental batch.

The Ozette Discovery platform can also be used to interrogate the distribution of marker expression across the identified phenotypes. For example, projection of CD3, CD20, or CD11b expression onto the Ozette UMAP shows that phenotypes identified by Ozette Discovery span all major PBMC cell lineages (Fig 2C). We can also examine the distribution of sample- or cell-level covariates. As is true for most biomedical experiments, the data published in Liu et al. had to be collected in several batches. This kind of batching potentially introduces unwanted variability in antibody staining and scRNA-seq sequencing results, and various algorithms have been proposed to correct for such batch effects^26^. However, because Ozette Discovery identifies cellular phenotypes on a per-sample basis and then combines annotations across samples, it intrinsically accommodates shifts in the protein expression distribution between samples, and therefore, batches. Correspondingly, Ozette Discovery phenotypes are well integrated across batches (Fig 2D).

Analysis of this dataset required a total of approximately 1.5 hours from preprocessed data to cell annotations, which compares favorably to the significant hands-on and computing time it often takes to batch correct, select markers, cluster, and annotate single-cell sequencing data using current state-of-the-art methods. We therefore find that we can use Ozette Discovery to annotate CITE-seq data, and that our method identifies granular, robust, and human-interpretable cell phenotypes.

### Ozette Discovery identifies more accurate phenotype labels than current methods

We next sought to compare how Ozette’s cell phenotype labels compare to the labels generated by Liu et al. using Louvain clustering, manual gating, and expert annotation. Liu and colleagues described 30 phenotypes across cells of the adaptive and innate immune systems, 23 of which (Fig 3A) were found in cells that passed our QC filters (see Methods). Overlaying these annotations onto the Ozette UMAP reveals that the authors’ phenotypes (colors) are distributed across multiple Ozette phenotypes (clusters), indicating that Ozette Discovery achieves greater resolution than canonical annotation methods (Fig 3B-D).

**Figure 3:**
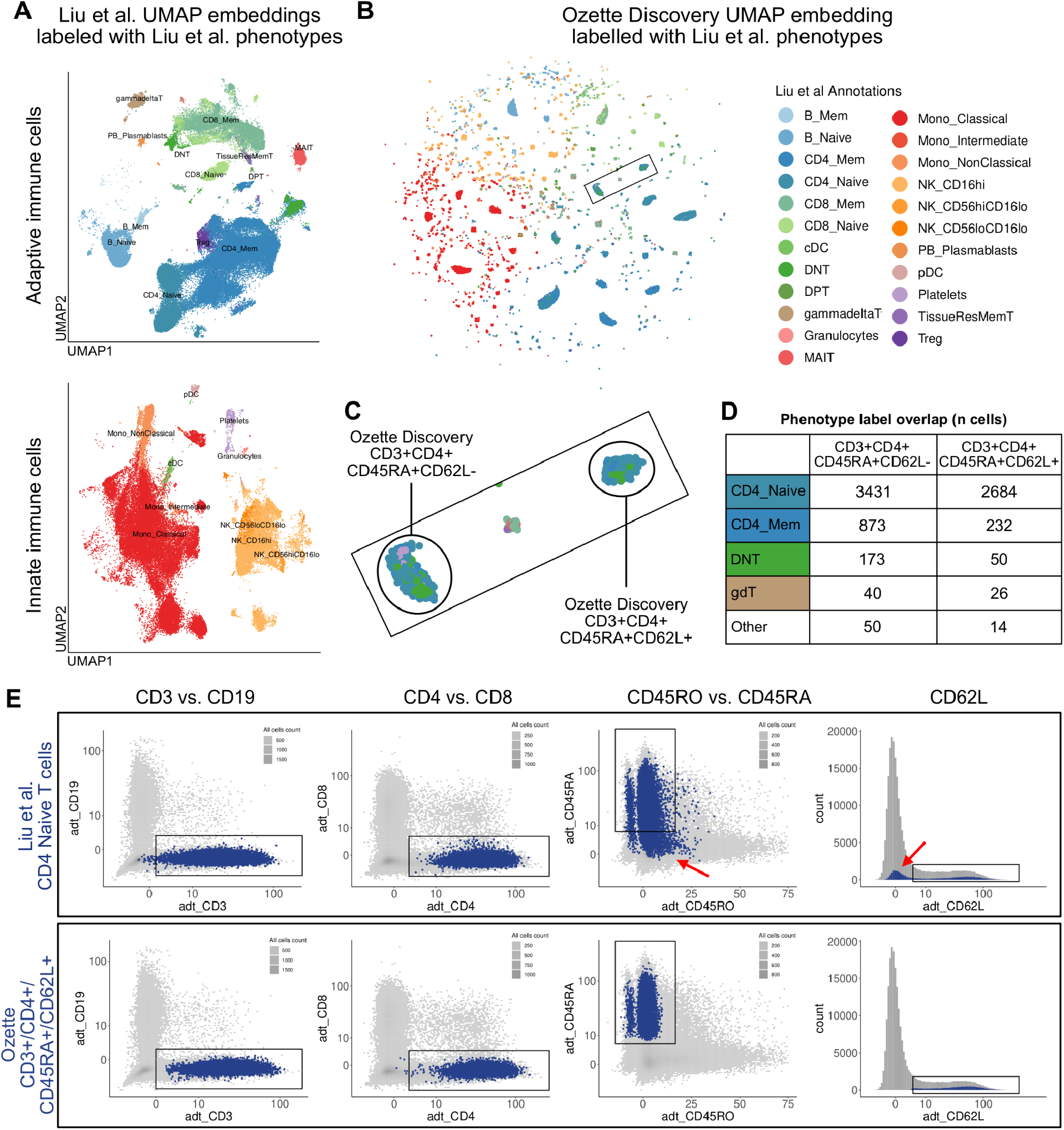
Phenotype labels generated by Ozette Discovery are more accurate than those generated by current CITE-seq annotation methods. **A)** Reproductions of UMAP embeddings presented in figure 2B of Liu et al., filtered to cells passing our QC metrics and colored by the authors’ coarse phenotype labels. Plots show cells of the adaptive immune system (top) and innate immune system (bottom). **B)** UMAP embedding generated by Ozette Discovery colored by Liu et al. phenotype labels presented in (A). **C)** Magnified inset from (B). Cells from multiple Liu et al. phenotypes (colors as in [B]) map to each Ozette phenotype (CD3+CD4+CD45RA+CD62L- or CD3+CD4+CD45RA+CD62L+). **D)** Table showing counts of cells annotated by Ozette Discovery as CD3+CD4+CD45RA+CD62L+ (left column) or CD3+CD4+Cd45RA+CD62L- (right column) versus their label in Liu et al. Liu et al. phenotypes that contributed less than 25 cells to either Ozette phenotype were grouped into an “other” category (9 phenotypes and 3 phenotypes, respectively). **E)** Backgating plots showing the expression patterns of CD4 naive T cells (blue) as defined by Liu et al. (top) or Ozette Discovery (bottom) compared to the distribution of all cells in the dataset (gray). Markers being plotted are indicated on the axes and at the top of the corresponding column. Black boxes indicate expected location of CD3+CD4+CD45RA+CD62L+ naive T cells, and red arrows indicate cells with aberrant ADT expression. Density of “all cells” is indicated by shades of gray. Blue color is not proportional to cell density. Scale is arcsinh transformed with cofactor = 7.

We also observed that cells from multiple different Liu et al. annotations fall into the same Ozette annotation (Fig 3B-D), indicating some discordance between the two labeling methods–Ozette Discovery is not simply further subdividing the authors’ annotations. To understand the root of this disagreement, we compared backgating plots showing the ADT expression profiles of matched cell types between the clusters provided in the original paper and Ozette Discovery’s annotations. For example, naive CD4 T cells are often defined in high-dimensional flow cytometry as CD4+CD45RO-CD45RA+CCR7+ ^27^. Liu et al. did not stain for CCR7, so we therefore substituted CD62L as a proxy^28^, yielding CD3+CD4+CD45RO-CD45RA+CD62L+ as the protein-based phenotype string for naive CD4 T cells. We find that naive CD4 T cells as annotated in Liu et al. (Fig 3E, top) sometimes include CD45RA- cells and CD62L- cells (Fig 3E, red arrows). In contrast, the Ozette phenotype annotated as CD19-CD3+CD4+CD8-CD45RO-CD45RA+CD62L+ (Fig 3E, bottom) by construction satisfies this sequence of gates. Similar irregularities were found with other cell types, including double negative T cells (Figure 7F). We therefore conclude that Ozette Discovery identifies more accurate and homogenous phenotypes than current unsupervised clustering methods.

Subsequently, we assessed how our ADT-based phenotype annotations compared with the transcriptomic profiles of the cells assigned to each phenotype. While our phenotypes are annotated exclusively based on protein expression data, a set of biologically meaningful phenotypes should also exhibit specific expression of expected canonical genes and pathways at the RNA level. Considering the most abundant phenotypes for the purposes of ensuring adequate statistical power, we generated pseudobulked gene expression data^29^ for each phenotype within each sample by adding up the raw unique molecular identifier (UMI) counts for each gene. We then performed differential gene expression analysis between each pair of phenotypes (see “Differential Expression Testing’’ in Methods). This analysis revealed that phenotypes of the same cell lineage as annotated by Ozette Discovery using the ADT data alone tend to also cluster together by gene expression (Fig 4A). Amongst the top differentially expressed genes between Ozette Discovery phenotypes, we found several canonical cell lineage markers including CD14, MS4A1, CD8A, CD3E, and NKG7 (Fig 4A, black arrows).

**Figure 4:**
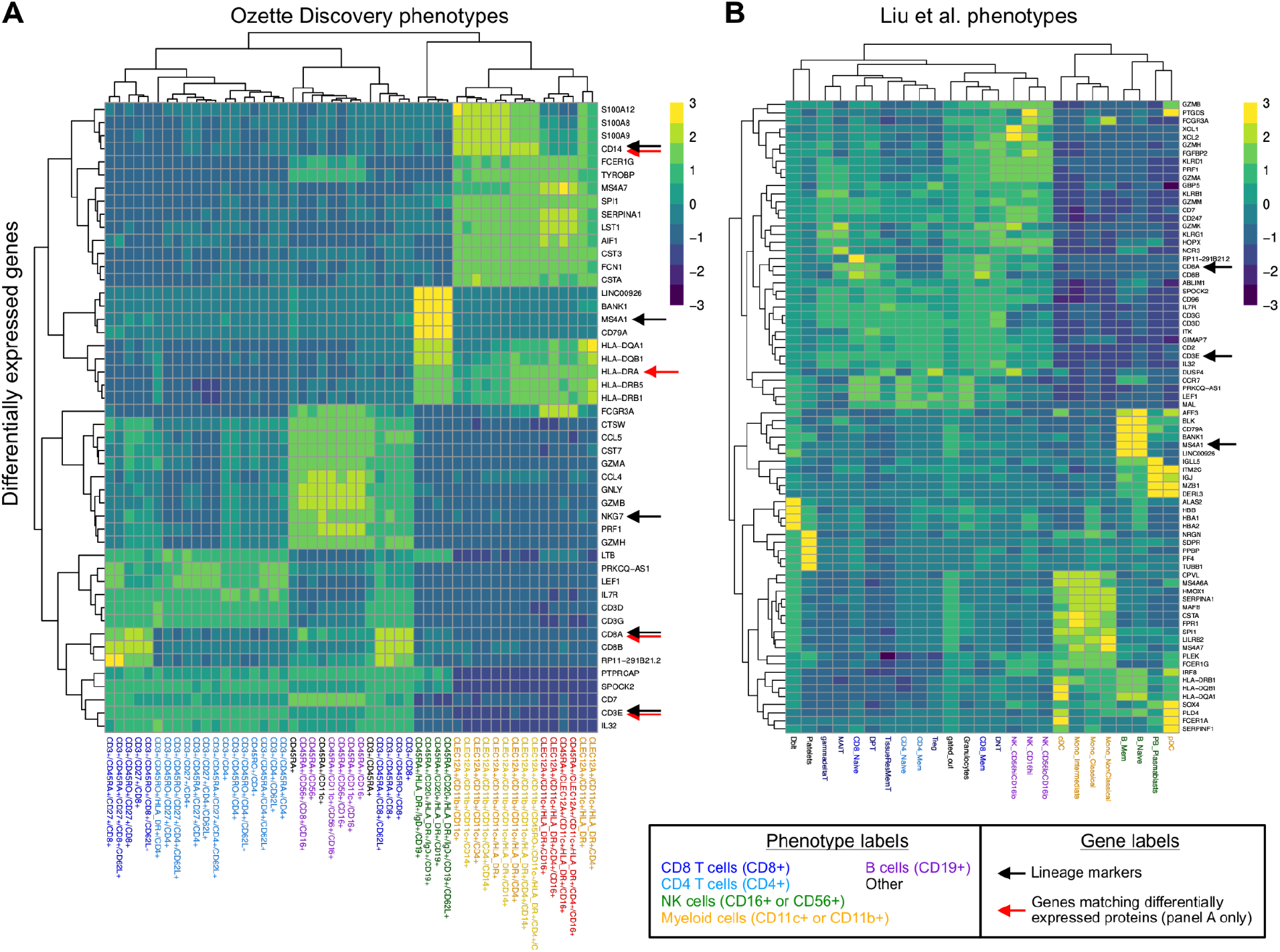
Phenotype labels generated by Ozette Discovery better correlate with RNA expression than current labeling methods. **A)** Heatmap showing up to the top 10 differentially expressed genes by logFC (rows) between cell phenotypes as defined by Ozette Discovery (columns). Color scale shows Z score of gene expression by row. Column names are colored according to cell lineage. Black arrows indicate canonical lineage markers and red arrows indicate genes encoding proteins that were used to determine Ozette Discovery phenotypes (legend in panel 4B). **B)** Heatmap showing up to the top 10 top differentially expressed genes by logFC (rows) between cell phenotypes as defined in Liu et al. (columns). Column names are colored according to cell lineage and black arrows indicate canonical lineage markers. In both panels, rows and columns of the heatmaps are ordered by hierarchical clustering of the euclidean distance between rows/columns using complete linkage.

Genes encoding for several of the ADT-measured proteins used to define our phenotypes were also identified as highly differentially expressed, including CD8A, CD3E, CD14, and HLA-D (Fig 4A, red arrows). Genes that are known to have discordant RNA and protein expression such as CD4^1^ were absent from the top differentially expressed genes (Fig 4A). When we performed the same analysis using the Liu et al. annotations, and found that phenotypes of the same cell lineage also tended to cluster together by gene expression, but the differences in gene expression between phenotypes and even cell lineages were less apparent than between Ozette’s annotations (Fig 4B). This analysis therefore suggests that despite annotating cell phenotypes using ADT protein expression alone, Ozette Discovery’s phenotype annotation method also captures many salient features of cells’ gene expression profiles.

### CLEC12A+CD11b+ cellular subsets are expanded in critical COVID-19 patients

The original study presented by Liu et al. identified several gene expression correlates of COVID-19 severity across moderate, severe, and critical COVID-19 cases (Fig 1A). The authors used the ADT data to annotate broad canonical cell types, and then focused on the gene expression data to identify correlates of disease severity. We hypothesized that the abundance of our ADT-defined phenotypes may also be associated with COVID-19 severity. To test this hypothesis, we performed differential abundance testing between moderate/severe and critical COVID-19 patients. To increase statistical power, we restricted our analysis to 109 sufficiently-abundant phenotypes and grouped all less-abundant phenotypes into a “rare” population. This analysis revealed a striking expansion of two cell populations in critical COVID-19 patients (Fig 5A): cells that were CLEC12A+CD11b+ or CLEC12A+CD11b+CD11c+ and negative for all other markers in the panel (Fig 5B-C). Moreover, the abundance of these phenotypes is also attenuated in healthy controls (Fig 5B), further indicating that this expansion is a specific hallmark of critical COVID-19 severity. While these cells express CD11b and appear to be monocytic, they lacked surface protein expression of CD14, though CD14 transcript was detectable (Fig 4A). Thus Ozette Discovery’s unbiased and exhaustive approach to cell annotation using surface protein expression can uncover novel phenotypic correlates of disease.

**Figure 5:**
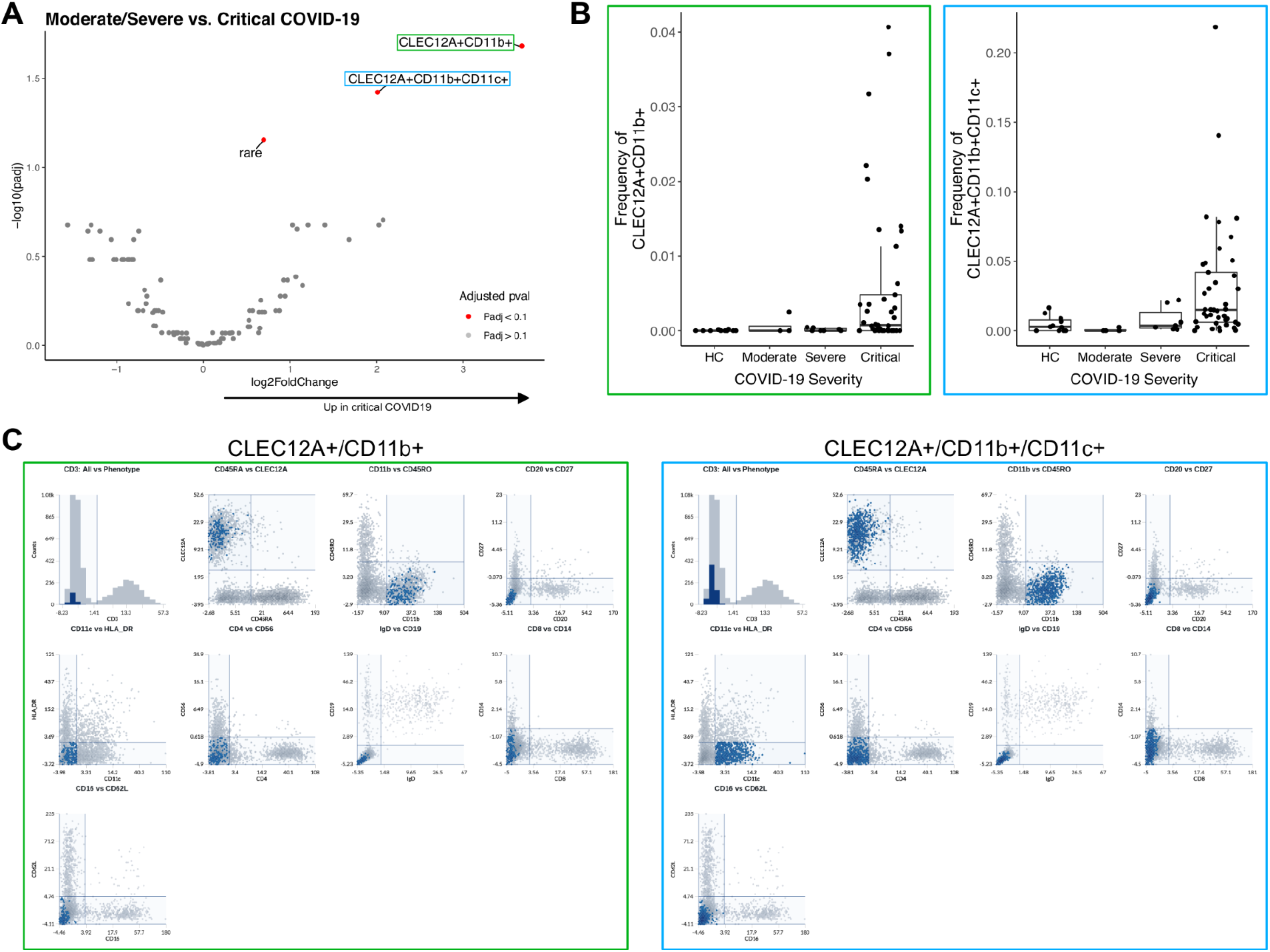
Identification of cell phenotypes expanded in critical COVID-19 patients. **A)**Volcano plot showing differential abundance of cell phenotypes, with phenotypes over-represented in patients with critical COVID-19 appearing to the right. Phenotypes with FDR q-value < 0.1 are highlighted in red and labeled. “Rare” indicates a catch-all group for low-abundance phenotypes (see Methods). **B)** Boxplots showing frequency (of total PBMCs) of phenotypes that are CLEC12A+CD11b+ or CLEC12A+CD11b+CD11c+ and negative for all other markers in the panel, stratified by COVID-19 severity. Dots represent individual samples, and boxes plot the median and first and third quartiles. Color outlining each plot matches the box outlining the location of that phenotype in (A). **C)** Backgating plots from the Ozette Discovery dashboard as described in Fig 2B, highlighting the two phenotypes boxed in (A). Color outlining each series of backgating plots matches the color outlining the corresponding phenotype in (A).

### Identification of RNA biomarkers that correlate with expansion of CLEC12A+CD11b+ cells

Since Liu et al. measured CITE-seq profiles longitudinally in COVID-19 donors, we next characterized the abundance of the most differentially abundant phenotype, CLEC12A+CD11b+, over time. Plotting the frequency of the CLEC12A+CD11b+ population versus days since symptom onset revealed that the expansion of this phenotype peaks around 20 days post-symptom onset (Fig 6A). Given this specific time period of expansion, we hypothesized that gene expression measured at time points prior to this expansion might be able to predict subsequent aberrant expansion of CLEC12A+CD11b+ cells. To explore this, we conducted differential expression analysis using data from the time point that preceded the maximum expansion of CLEC12A+CD11b+ cells for each COVID-19 patient. We first performed this analysis in an All Cells pseudobulk, where we summed the gene expression of every cell in every Ozette phenotype within each sample. Using this All Cells pseudobulk approach we found over 400 genes to be differentially expressed (a subset of most significantly differentially expressed genes is shown in Fig 6B, top row), with many related to interferon signaling (Fig 6B). This is consistent with previous work demonstrating that changes in interferon signaling are associated with poor COVID-19 outcomes^30,31^. We also found that early upregulation of IL2RA (encoding CD25 protein) was associated with expansion of CLEC12A+CD11b+ cells (Fig 6C). This is consistent with Liu et al.’s finding that CD25 protein levels were elevated in circulating blood of critical COVID-19 patients at early time points in a separate COVID-19 cohort (see Liu et al. Fig 6B). We then repeated this analysis, generating pseudobulked profiles within each Ozette Discovery phenotype, in order to test for cell type-specific predictors of CLEC12A+CD11b+ expansion. This analysis again highlighted IL2RA, which was significantly upregulated in CD3+CD45RO+CD4+CD62L+ and CD3+CD45RO+CD4+CD62L- cells (FDR < 0.03 and FDR < 0.09, respectively), and upregulated (log fold change > 3) but not at FDR-significant levels in five other CD3+CD4+ subsets. We additionally identified genes that were only significantly up- or down-regulated within specific lineages and not in the All Cells pseudobulk, such as interferon-inducible IFNGR2 (upregulated specifically in CD3+/CD45RA+/CD27+/CD4+ cells) and IFI44 (upregulated specifically in CLEC12A+CD11b/CD11c+/CD4+/CD14+ cells) (Fig 6B-C). This suggests that interferon signaling specifically within other cell types may be driving the expansion of this CLEC12A+CD11b+ phenotype in critical COVID-19 patients. We therefore conclude that phenotypes identified by Ozette Discovery elucidate disease etiology by identifying gene expression predictors of cell type abundance.

**Figure 6:**
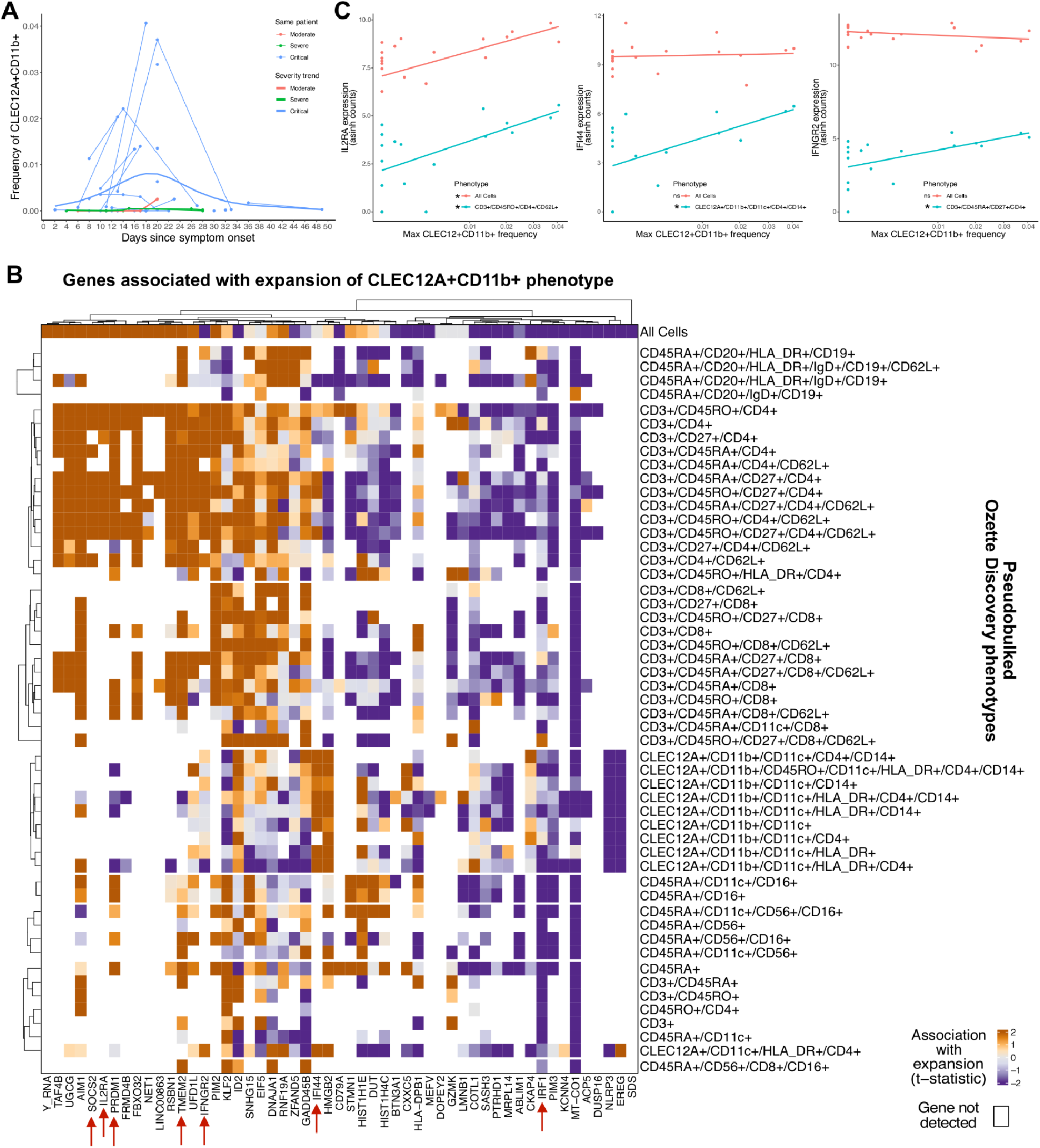
Changes in interferon-related gene expression precede expansion of CLEC12A+CD11b+ phenotype. **A)** Days since COVID-19 symptom onset versus frequency (of total PBMC) of CLEC12A+CD11b+ cells. Each point represents a sample, and thin lines connect samples from the same patient. Thick lines indicate average trend by COVID-19 severity. Dots and lines are colored by COVID-19 severity. **B)** Heatmap showing genes that are differentially expressed in accordance with expansion of the CLEC12A+CD11b+ phenotype. The 10 most significant genes by p-value and FDR < 10% within each Ozette Discovery phenotype are shown. Empty cells represent tests that were not possible because that gene was not detected in the corresponding phenotype. Genes shown in panel (C) are indicated with red arrows. **C)** Plots showing expression of selected genes versus maximum frequency of CLEC12A+CD11b+ phenotype in either all cells or the indicated Ozette Discovery phenotype. ns = not significant. *= significant at FDR < 10%.

### Identification of early RNA biomarkers of COVID-19 severity

Based on the results of this approach, we next attempted to use the gene expression data from the earliest time points sampled for each patient to identify mRNA predictors of COVID-19 severity, without regard to the expansion of any cell types. To determine the relative utility of our Ozette Discovery phenotypes, we performed this analysis in an All Cells pseudobulk (L1), cell lineage pseudobulk (L2), and at the level of each individual Ozette Discovery phenotype (L3) (Fig 7A). We found that no genes were differentially expressed in the All Cells pseudobulk (L1), and only 3 genes within the CD4 T cell lineage were differentially expressed at the L2 level (Fig 7B, left). Yet when we repeated this approach for the most granular Ozette Discovery phenotypes, we found 131 genes differentially expressed at 10% FDR (Fig 7B, left). Strikingly, among these 131 genes differentially expressed at early time points, we identified increased GZMB and PRF1 gene expression in CD3+CD45RA+CD4+CD62L+ naive T cells from patients with higher COVID-19 severity (Fig 7C, black asterisks). Critical patients expressed 5-fold higher GZMB and 2.8-fold higher PRF1 transcript levels than severe patients (Fig 7D, top). This higher level of GZMB and PRF1 gene expression was conserved across several distinct CD4 T cell populations, but was only detectable at conventional FDR cutoffs in CD3+CD45RA+CD4+CD62L+ naive T cells (Fig 7C, red box). Upregulation of GZMB and PRF1 is a hallmark of cytotoxic function, and so this gene expression pattern within putative naive CD4 T cells may signal the early transition of these CD3+CD45RA+CD4+CD62L+ naive T cells towards a cytotoxic state^32^.

**Figure 7:**
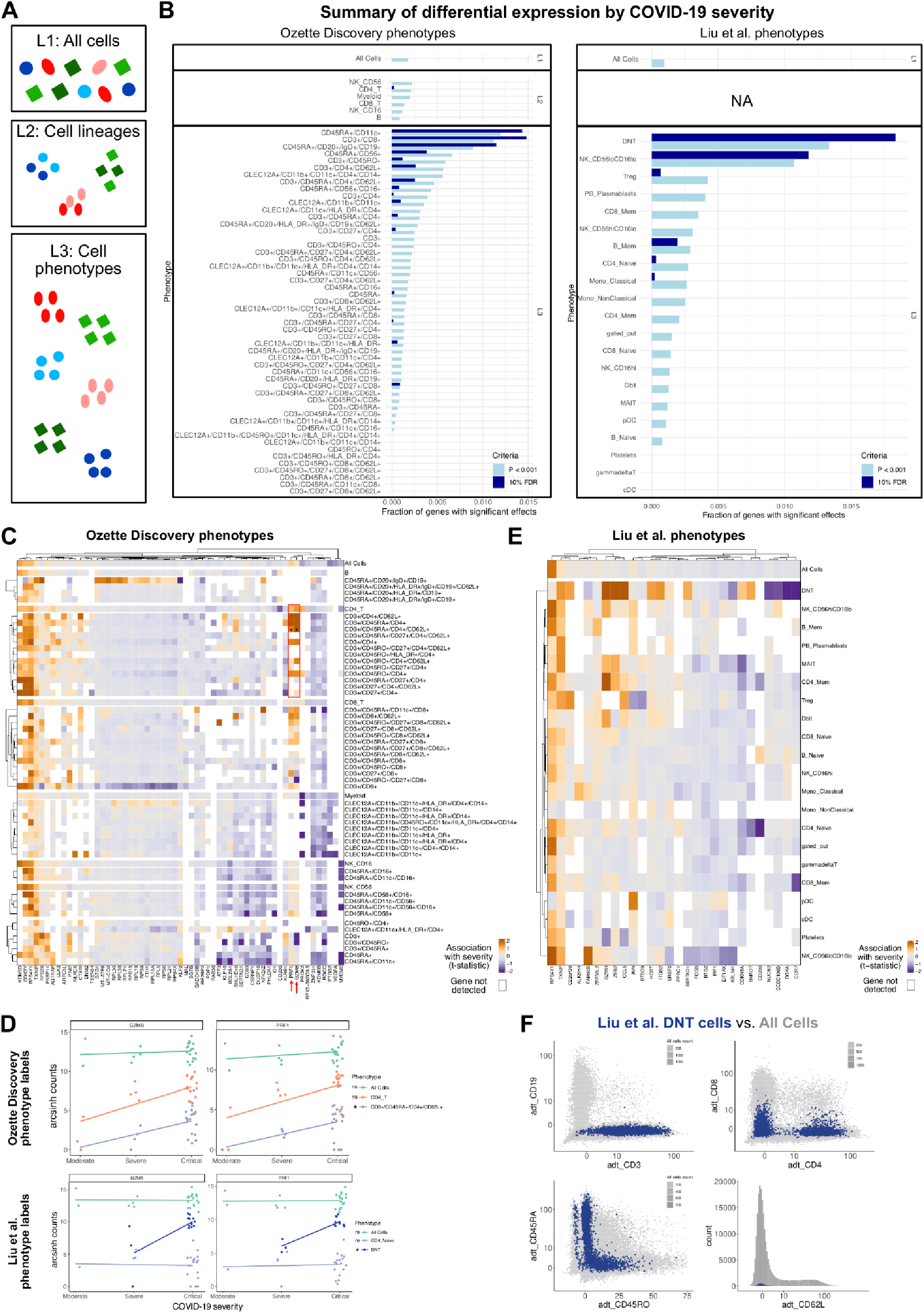
Identification of early predictors of severity. **A)** Schematic illustrating pseudobulking approach used in this figure: gene expression for each cell (represented by colored shapes) is aggregated by pseudobulking across phenotypes into L1 (all cells; coarsest), L2 (cell lineages; intermediate), or L3 (preserving Ozette Discovery phenotypes; most granular) levels to perform differential expression testing. **B)** Fraction of genes tested that are differentially expressed between COVID-19 severity levels at early time points at either p < 0.001 (light blue bars) or FDR < 10% (dark blue bars) in the indicated populations. Left panel shows this analysis using Ozette Discovery phenotype labels and the right panel shows the same analysis using the cell phenotype labels provided in Liu et al. **C)** Heatmaps showing the top differentially expressed genes for L1, L2, or L3 Ozette Discovery phenotypes. Formatting is the same as for Fig 6B. Stars indicate significant upregulation of GZMB and PRF1 in CD3+CD45RA+CD4+CD62L+ phenotype Fig 2D. Red arrows and red box indicate t-statistics for GZMB and PRF1 differential expression. **D)** Plots showing expression of GZMB and PRF1 by COVID-19 severity in the indicated Ozette Discovery phenotypes (top) or Liu et al. phenotypes (bottom). ns = not significant. * = significant at FDR < 10%. **E)** Heatmap as in panel (C) showing L1 “All Cells” and L3 Liu et al. phenotypes. **F)** Backgating plots showing ADT expression of cells labeled as DNT by Liu et al. (blue) compared to all cells (gray). Formatting is the same as in Fig 2B.

We then repeated this analysis using the authors’ annotations as the L3 classifications. Again, we found no genes differentially expressed at 10% FDR in the All Cells pseudobulk, but we did find a total of 91 differentially expressed genes across the authors’ phenotypes (Fig 7B, right; 7E). This list also included GZMB and PRF1, but only within “DNT’’ cells, which the authors do not define in their original publication but likely refers to double negative T cells^33^ (Fig 7D, bottom). However, we assessed ADT expression patterns of the authors’ DNT population using backgating plots and found that this population had varied expression of CD4, CD8, CD45RA, CD45RO, and CD62L (Fig 7E). Furthermore, we found that 15% of the DNT phenotype was classified by Ozette Discovery as CD3+CD4+CD45RA+CD62L± cells (data not shown), consistent with these phenotypes showing an upregulation of GZMB1 and PRF1 with increasing COVID-19 severity. Interestingly, the DNT subset does form two distinct clusters adjacent to memory CD4 T and memory CD8 T cells in the authors’ UMAPs (Fig. 3A; Liu et al Fig 2B) suggesting that this DNT phenotype may represent an early shift towards effector function in a mixture of T cell subsets, as opposed to a cell type that could be easily and reproducibly isolated via flow cytometry. We therefore find that phenotypes identified by Ozette Discovery can be used to identify cell type-specific predictors of disease severity.

## CONCLUSIONS

Here we demonstrated that Ozette Discovery, which was developed to provide comprehensive and interpretable cell phenotype annotations using protein expression in cytometry data, can be applied to sequencing-based protein expression data derived from technologies such as CITE-seq. Our pipeline is compatible with well-established preprocessing methods currently used for single-cell sequencing data, but the addition of our novel background normalization method significantly improves the signal-to-noise ratio and therefore the utility of ADT data. Ozette Discovery also intrinsically accommodates shifts in location or scale of ADT protein expression distributions, resulting in phenotypes that are robust to batch effects without the need for additional batch correction methods. While both current methods and Ozette Discovery can provide exhaustive annotations, Ozette Discovery generates phenotype labels that are more human-interpretable in terms of their names and derivations. The net result is a set of phenotype labels that are more granular, robust, homogeneous, and human-interpretable in a fraction of the time, compared to current state-of-the-art annotation methods.

We harnessed the richness of these phenotypes by quantifying their abundance to identify correlates of disease. While this is a common approach in scRNA-seq studies, it is rarely dispositive, since the phenotypes are not sufficiently granular or interpretable. This fundamental analysis revealed two groups of CLEC12A+CD11b+CD14- myeloid cells that were expanded specifically in critical COVID-19 patients. The canonical monocyte marker CD14 was not detected in these phenotypes at the protein level, despite the fact that the CD14 transcript was detected (Fig 4A). Furthermore, the CLEC12A+CD11b+CD14+ populations were not significantly associated with critical COVID-19. This indicates that absence of CD14 protein expression is a key feature of the CLEC12A+CD11b+ populations associated with critical COVID-19, underscoring the importance of interrogating both the gene and protein expression patterns of cell phenotypes.

That is not to say that interrogation of the RNA expression patterns is not a worthwhile analysis–the homogeneity within Ozette Discovery’s phenotype labels provide a solid foundation for interrogating differential gene expression. This homogeneity also means that simple and robust methods, such as pseudobulking, are appropriate to use on these data for differential expression testing. Indeed, we were able to use pseduobulking within Ozette Discovery phenotypes to unbiasedly elucidate early transcriptomic patterns associated with the expansion of CLEC12A+CD11b+ cells, including increased expression of IL2RA (CD25) and interferon-inducible genes.

This pseudobulking approach also successfully identified early correlates of subsequent COVID-19 severity, regardless of expansion of CLEC12A+CD11b+ cells. While there were no genes differentially expressed across pseudobulks comprised of all cells, and very few genes differentially expressed at the level of cell lineage, PRF1 and GZMB were positively associated with COVID-19 severity in a specific subset of naive CD4 T cells. PRF1 and GZMB are genes associated with T cell effector capability, and so this finding may indicate that this subset of naive T cells are starting to transition towards a cytotoxic state. This therefore highlights how our approach fully exploits the strengths of multi-modal data: identifying a homogeneous cell phenotype using the ADT protein expression data, and then identifying early gene expression changes within that cell type that may indicate its future commitment. We also note that because our phenotype labels are not derived from gene expression data, such follow-on investigations are far less susceptible to p-value distortion due to the “double dipping” that occurs when gene expression data is used both to derive the cluster identity and to test for differential expression^34^. Our analysis pipeline therefore fully harnesses the strength of the CITE-seq assay by using Ozette Discovery and the ADT data to identify differentially abundant phenotypes, and then using the paired gene expression data to identify potential mechanisms for this change.

The difficulty of translating *in silico* findings from sequencing experiments into follow-up experiments has been widely recognized: often the results from a sequencing experiment fail to replicate using alternative methods like flow cytometry. CITE-seq assays offer progress, as they have the potential to synergize with existing protein-based cell annotations used in cytometry. However, without careful analysis of the ADT data, one can still be led astray. Both Liu et al. and our annotations identified specific over-expression of PRF1 and GZMB in subsets of cells at early time points in critical COVID-19 patients. In Liu’s annotations, this over-expression was attributed to “DNT” cells, which in our analysis were not generally double negative, but rather a mixture of CD4+, CD8+ and CD4-CD8- T cells. It is possible that differences in data transformation contribute to this discordance: we employed the novel background normalization technique described here, in contrast to Liu et al.’s use of DSB^23^. These preprocessing differences may alter the threshold locations between positive and negative populations, and therefore produce discordant phenotype labels. Nevertheless, the DNT cells as annotated by Liu et al. lacked an obvious pattern of protein expression in common lineage markers that would allow them to be reproducibly isolated and studied. In contrast, we show that the annotations derived from Ozette Discovery are homogeneous and definitive, and therefore readily lend themselves to the design of a follow-up flow cytometry study.

Our study of differential expression in the Liu et al. dataset therefore suggests that a narrow focus on gene expression in CITE-seq data poses the risk of mis-classifiying cell types and thereby mis-attributing effects of interest. In contrast, protein-based annotations can leverage decades of domain knowledge about the characteristics of various lineages. However, the cell phenotypes discoverable by our method are limited to the markers included for analysis. We credit that some antibodies commonly used in flow cytometry, such as CCR7 and antibodies for gamma-delta T cells, remain difficult to stain for in assays such as CITE-seq at sufficient concentrations to allow clear thresholds to be drawn without contaminating the sequencing pool with unbound ADT. We also credit that some cellular states, for example cell cycle phase, are also often easier to identify via gene expression than protein expression. In these cases, the mRNA can serve as a proxy for protein expression, and we expect that additional methodology will soon be developed to allow this.

In total, our work illustrates that Ozette Discovery provides a rich and meaningful set of cell phenotypes, enabling scientists to harness the advantages of both cytometry and single-cell sequencing at once.

## INQUIRIES

We welcome correspondence to.

## METHODS

### Data curation

Unfiltered HDF5 files were downloaded from GEO GSE161918. The original study included both PBMC-derived cells as well as a smaller fraction of cells enriched for non-naive B and T cells by flow sorting. We excluded the sorted cells from this analysis due to staining artifacts, possibly due to competition between the ADT antibodies and flow sorting antibodies. R 4.1.3 and DropletUtils 1.19.1 were used to initially identify potential cell-containing droplets. Hashtags were resolved using demuxmix version 1.2.0, and candidate doublets identified with scDblFinder 1.8.0. Ambient background normalization for ADT expression was performed using custom-developed methods (see “Background normalization” below). After QC metrics on the number of genes detected, empty droplet probability, mitochondrial fraction, doublet probability, ADT fraction, and minimum cell count per sample were applied, 213,679 cells remained. These cells were derived from 63 samples across 43 donors (n = 11 healthy controls, n = 3 moderate COVID-19 patients, n = 5 severe COVID-19 patients, and n = 24 critical COVID-19 patients). For the purpose of depicting cell annotations published in Liu et al. (Fig 3A, B, D, E, Fig 4B, Fig 7D, E, F), we used the phenotypes labeled as “WCTcourse” in the source code of Liu et al.

### Background normalization

In each 10X lane, we suppose that ADT counts for each barcode and feature arise as a sum of an ambient source, plus other (i.e. cell-bound) sources. The ADT counts in non-cell-containing droplets are assumed to arise from only ambient sources, and are used to estimate the relative abundance vector **a**=[*a*_i_] of each feature *i* in the ambient pool. In each cell-containing droplet, we decompose the total sum of ADT UMIs across features into an ambient sum *N_a_*, and residual sum *N_r_* by regressing the observed UMI counts onto the ambient relative abundance vector **a**. Then, for each cell-containing droplet and each feature *i* we estimate the ambient contribution in a generalized linear model using *a_i_* as a covariate and log(N_a_) as an offset. The Pearson residuals from this regression form the background-corrected expression values.

### Clustering

Background-corrected ADT expression of 17 markers was used to annotate cell types using the Ozette Discovery platform version 4.5, and the phenotypic identity of each cell was imported into R. 170 phenotypes were discovered. The Ozette UMAP (Fig 2A, C, D, Fig 3B) employs an annotation transformation step (https://github.com/flekschas-ozette/ismb-biovis-2022) before calculating the embedding. For each expression category in a marker (i.e., negative and positive categories), the level-specific marker expressions are winsorized to remove outliers, normalized (in each sample) to have zero mean and unit variance, and shifted to category-specific location to ensure co-localization of cells from the same phenotype.

### Differential Abundance Testing

Phenotypes present in at least 5 samples and with a median of 1 cell across all samples (109 phenotypes) were converted into a phenotype-samples count matrix, which was used as input into DESeq2^35^ version 1.34.0. Cells belonging to phenotypes that did not meet these criteria were aggregated into a “rare” catch-all category to preserve the total number of cells in each sample, i.e. for calculating size factors. We fit a model comparing severity, encoded as a factor (encoded as healthy, moderate/severe, critical) and adjusting for a technical batching variable as provided in the original study. To estimate the abundance of the CLEC12A+CD11b+ cells across time in Fig. 6A, a negative binomial gam was estimated with mgcv version 1.8.33 ^36^. The log total number of cells was provided as an offset and the resulting predicted counts were converted into proportions.

### Differential Expression Testing

Differential expression tests were in all cases conducted using pseudobulked gene expression profiles. The UMI counts of each gene over each sample and phenotype were summed to yield the L3 pseudobulked expression. Here, we required phenotypes to be present in at least 1 cell in 5 samples, and 7 cells in the median sample. The L2 and L1 count matrices were formed analogously by summing over combinations of phenotypes that were annotated as expressing combinations of key lineage markers (CD3, CD4, CD8, CD11b, CD16, CD19, CD56), or all cells, respectively.

To test for differential expression between phenotype labels (Fig 4), we normalized the pseudobulk profile with its library-size, transformed with the hyperbolic arcsin using `scuttle::logNormCounts(…, transform = “asinh”)` and then found differentially expressed genes via t-tests in `scran::findMarkers.` We considered the 5 genes with the most significant p-values, as long as that gene had FDR < 0.1.

For the associations with CLEC12A+CD11b+ abundance, for each COVID-19 positive donor sampled no later than 20 days since symptom onset (n = 23), we took a cross-sectional snapshot of the data by taking the sample from the time point preceding maximum abundance of CLEC12A+CD11b+ cells in that donor, or if that occurred in the first time point, then we took the first time point. We employed a square-root transformation of the CLEC12A+CD11b+ frequency as the independent variable in the regression and adjusted for technical batch. We tested for differential expression within each L3 and L1 phenotype using the function `pseudobulkDGE` with `method=”voom”` in the package `scran` version 1.22.1, and reported up to 10 genes per phenotype that had FDR < 0.1 in the presented heatmap.

For the associations with severity, we encoded severity as an integer, with 2 indicating moderate, 3 indicating severe and 4 indicating critical and used this score as a quantitative variable, again adjusting for technical batch. We derived a cross-sectional snapshot by taking the first time point for each donor, as long as that time point occurred before 20 days post-symptom onset. If no samples occurred before 20 days post-symptom onset, then we omitted that donor. We tested for differential expression within each L3, L2, and L1 phenotype using the function `pseudobulkDGE` with `method=”voom”` in the package `scran` version 1.22.1, and reported up to 10 genes per phenotype that had FDR < 0.1.

